# Improving Statistical Rigor in Animal Aging Research by Addressing Clustering and Nesting Effects: Illustration with the National Institute on Aging’s Intervention Testing Program Data

**DOI:** 10.1101/2025.03.14.642436

**Authors:** Erik S. Parker, Lilian Golzarri-Arroyo, Stephanie Dickinson, Beate Henschel, Luis-Enrique Becerra-Garcia, Thirupathi R. Mokalla, Olivia C. Robertson, Deependra K. Thapa, Colby J. Vorland, David B. Allison

## Abstract

Clustering effects, such as those introduced by housing animals in shared cages, are often overlooked in preclinical lifespan studies, despite their potential to distort variance estimates and inflate Type I error rates, leading to misleading conclusions. This methodological oversight reduces statistical rigor and may undermine the reliability of findings. To address this gap, the current study examines the impact of accounting for clustering and nesting effects on lifespan analyses by comparing the results of statistical models which both account for and ignore these effects. Using 2019 data from the Interventions Testing Program (ITP), a large-scale initiative evaluating the effects of compounds on lifespan in UM-HET3 mice as a case study, we illustrate how different modeling approaches influence statistical estimates and conclusions.

Clustering and nesting effects were addressed using linear mixed effects, and Cox frailty models, both of which explicitly account for cage-level dependencies and different levels of data nesting. Comparisons were made between unadjusted lifespan analyses and those incorporating clustering and nesting adjustments. The results of this case study indicate that properly adjusting for clustering and nesting effects can change the conclusions drawn from statistical significance tests as compared to unadjusted model approaches, and so it remains best practice to properly account for clustering and nesting to reduce the potential for inflated Type I error rates.

These findings highlight the importance of accounting for clustering and nesting in preclinical research to ensure valid and robust statistical inference. By demonstrating the practical application of clustering adjustments, this work underscores the broader implications for improving reproducibility and rigor in lifespan studies and other experimental designs.

## Introduction

In aging research, preclinical studies often rely on animal models to evaluate the effects of interventions on lifespan and healthspan. A persistent methodological challenge in these studies is the failure to account for clustering and nesting effects inherent in experimental designs. Clustering arises when observations within a group or shared environment, such as animals housed in the same cage, may exhibit correlated outcomes. Nesting refers to a hierarchical data structure where clusters (e.g. cages) are nested within treatment, resulting from a cluster randomized controlled trial (cRCT). When clustering in the data is ignored in analysis, standard errors of estimates may be artificially reduced, such that there is an increased risk of Type I errors (**Ball & Rabe, 2020; Li & Redden, 2015; Murray et al., 2004**; **Chusyd et al., 2022; Golzarri-Arroyo et al., 2022**; **Jamshidi-Naeini et al., 2022**). Similarly, false positives (Type I error) can result from using degrees of freedom at the level of the individual animal rather than the number of cages in the context of nesting from a cRCT (**Ball & Rabe, 2020; McDermott et al., 1994; Leyrat et al., 2018; Li & Redden, 2015**). Despite decades of methodological guidance (**Festing and Altman, 2002**; **McDermott et al., 1994**), clustering and nesting effects are still frequently overlooked in preclinical studies, including those in aging research, and more must be done to account for these effects to achieve valid statistical inference (**Brown et al., 2015)**.

Fortunately, statistical methods such as linear mixed models, generalized estimating equations, and Cox frailty models provide researchers with robust tools to account for clustering and nesting effects using widely available software packages such as R and SAS (**Bates et al., 2015; Kahan et al., 2016**; **Leyrat et al., 2018**). These methods extend standard approaches by modeling dependencies within clusters, ensuring that statistical conclusions accurately reflect the true structure of the data. However, as in other fields, we observe that researchers in the aging field often fail to account for clustering and nesting, underscoring the need for greater statistical rigor.

Our paper illustrates the concepts and methods to account for clustering and nesting with a case study by analyzing data from the Interventions Testing Program (ITP), a multi-site initiative investigating the effects of various compounds on lifespan in genetically heterogeneous UM-HET3 mice. Data from a recent ITP study **(Harrison et al., 2024)** are analyzed for illustration. We applied statistical methods that both ignore and account for clustering and nesting. All code used for the analyses is shared publicly to promote transparency and facilitate ease of implementation for researchers.

## Methods

### Accounting for Clustering and Nesting in Statistical Analyses, Generally

When analyzing data with inherent clustering or nesting, it is crucial to account for these structures to ensure accurate statistical inference. Several established methods address how to account for clustering effects, including general, and generalized linear mixed models (GLMMs), which incorporate random effects to model the dependencies within clusters (**Bates et al., 2015, Murray et al., 2004; Raudenbush & Bryk, 2002; Snijders & Bosker, 2012**), and generalized estimating equations (GEEs), which provide robust standard errors by accounting for within-cluster correlations at sufficiently large sample sizes (**Hardin & Hilbe, 2003; Li & Redden, 2015**). For time-to-event data, Cox frailty models extend the Cox proportional hazards (PH) model by incorporating random effects (frailties) to account for unobserved heterogeneity within clusters (**Therneau & Grambsch, 2003**). In addition to accounting for clustering with random effects, it is also vital to account for nesting in a cRCT. When the clusters (e.g. cages) are nested under each treatment, as in a cRCT, the degrees of freedom should be based on the number of cages rather than the number of individuals, as the cages are the independent unit of analysis. Kenward-Roger estimation is one method which provides appropriate degrees of freedom (df) which have been shown to maintain the Type I error rates in analysis. The impact of degrees of freedom adjustment method is highest in designs with a low (n < 30-40) numbers of clusters (**Kahan et al., 2016; Leyrat et al., 2018; Li & Redden, 2015**). Using SAS PROC MIXED, nesting can be specified in the random statement e.g. “random intercept/subject=cage_id (trt)” and the Kenward-Roger method will appropriately adjust the df for the model when *ddfm = kenwardroger* is specified on a random statement e.g. “random intercept/subject=cage_id”. In R, the df can be adjusted for nesting using the Kenward-Roger method by using the *ddf* = *“Kenward-Roger”* and *lmer*.*df = “Kenward-Roger”* options in the *anova* (when the mixed model was fit with the *lmerTest* function) and *emmeans* functions.

### Case study using ITP survival data

The 2019 ITP data used in this analysis was generated from an experiment testing the effects of multiple compounds on lifespan in UM-HET3 mice housed at three test sites (University of Michigan, University of Texas, and the Jackson Laboratory) **(Harrison et al., 2024)**. Briefly, mice were weaned at 42 days of age and assigned to a cage, after which cages were then randomized using a block design. All mice within the same cage were given the same treatment (as it was included in their food), and randomization was done separately for male and female animals at each test site. There were 777 total cages spread across the three test sites. The treatments used in this study (in addition to the control group) were Astaxanthin (*Asta*), Meclizine (*Mec*), Dimethyl fumarate (*DMF*), Mycophenolic acid (*MPA*), 4-Phenylbutyrate (*PBA*), and *SG1002* – a hydrogen sulfide donor. Full information on the mice rearing methods can be found in **Miller et al., 2011** and **Harrison et al., 2024**.

Below we outline our use of linear mixed and Cox frailty models, as applied to these ITP survival data as a case study.

### Analysis not Accounting for Clustering and Nesting

We first analyzed the ITP data with standard approaches for lifespan analysis including log-rank tests and Wang-Allison ‘maximum’ lifespan tests (tests of upper quantiles of the survival distribution, not true maxima), separated by sex, for analysis of each treatment group versus controls, without accounting for clustering and nesting. The log-rank tests were stratified by site, and the Wang-Allison tests were performed exclusively on non-censored animals who had observed deaths recorded. Cox proportional hazards (PH) models were performed, unadjusted for cage, by running separate models for each group versus controls, to demonstrate that a Cox model unadjusted for covariates is equivalent to the log-rank test (**Stefanescu, 2003**; **Harrell, 2015)**. Further, we also fit “full” linear models where the number of days alive was treated as a continuous outcome (on only the non-censored animals) and Cox PH models that included all treatment groups and both sexes with interactions between sex and treatment, unadjusted for cage effects.

### Analysis Accounting for Clustering and Nesting

To assess the impact of cage clustering effects, we first fit linear mixed models, with random intercepts for cage nested within treatment where the number of days alive was treated as a continuous outcome for all non-censored animals (because very few animals in this study were censored, 66 out of 2635 total animals). We chose to use this approach because when there is no censoring (i.e., missing survival information) of subjects, linear mixed models have favorable properties that have been well studied – though this approach is theoretically less powerful than the Cox frailty models (**Austin, 2017**). Because the linear (mixed) model framework is likely more familiar to most researchers than the Cox frailty model, we include both methods to illustrate the similarity to the results that can be obtained using the more powerful Cox frailty approach.

Our linear mixed models were fit with all treatments and sexes included. In these “full” models, a categorical treatment variable interacted with categorical sex and the main effects of both variables were included as fixed effects. A “full” model including all data was fit here primarily because including all the data in a single model increases the statistical power of the analysis as it uses the entire dataset to more accurately estimate the effects of the variables in the model, thus leading to more precise estimates of treatment effects (**Ouyang et al., 2020**). In our linear mixed model, cage nested within treatment was included as a random intercept to capture the degree to which animals housed in the same cage might have survival times more similar to each other than to animals housed in some other cage, due to “random”, unmeasured factors that are not accounted for in the model. Denominator degrees of freedom were adjusted using the Kenward-Roger approximation to be at the level of the number of cages rather than individual animals. Statistical significance of the interaction term and main effects were assessed using type 3 sum of squares, and estimated marginal means were compared to assess significant differences in lifespan between each treatment group and the control group within each sex.

Finally, we fit similar “full” Cox frailty models which similarly included a categorial variable for all treatments interacting with a categorical variable for sex. The frailty model is an extension of the standard Cox PH model which incorporates random effects (the frailty terms), much like the linear mixed model, to account for dependence, correlations, or unobserved relationships within the data (**Balan & Putter, 2020**). These frailty models were fit using both gamma and Gaussian distributions for the random effect frailty terms of cage nested within treatment. The gamma and Gaussian distributions are both commonly used but have different mathematical characteristics and may perform better in certain scenarios (**Balan & Putter, 2020**). Because we did not have an a priori reason to believe one would perform better than the other, we used both distributions here and compared the results of the two. Degrees of freedom for the Cox frailty models are estimated following **Therneau & Grambsch, 2003**. Type 3 sums of squares were used to assess the significance of the interaction term and main effects, and estimated marginal means were calculated and compared to assess significant differences in death hazards between each treatment group and the control group within each sex.

Log-rank and Wang-Allison tests were performed in R (version 4.3.0) using the *survival* package (version 3.5-5; **Therneau and Lumley, 2015**). Linear mixed models and Cox frailty models were fit in both SAS (version 9.4) and R using the *survival* (version 3.8-3; **Therneau & Lumley, 2015**), *lme4* (version 1.1-35.1; **Bates et al., 2015**), and *emmeans* (version 1.8.8; **Russell, 2018**).

## Results

### Standard Approaches not Accounting for Clustering or Nesting

When applying the standard analytic approach of pairwise log-rank/Cox-models unadjusted for cage effects nor accounting for nesting within site, we found that male mice treated with *Asta* and *MEC* had significantly longer median survival time as compared to the control group (908 vs 814 days, *p* = 0.003; and 881.5 vs 814 days, *p* = 0.027; Table 1a). Animals in the *Asta* and *MPA* were seen to have significantly longer “maximum” lifespan as per the Wang-Allison test (1144 vs 1079 days, *p* = 0.022; 1139.5 vs 1079 days *p* = 0.050; Table 1a).

**Table 1a.** Standard lifespan data analyses, unadjusted for cage - for male animals only. Note that we have included p-values from both the log-rank test and unadjusted Cox models to illustrate that they are equivalent tests when unadjusted for random effects or additional covariates (*Stefanescu, 2003;* Harrell & Harrell, 2015).

|  |  |  |  | Males |  |  |  |  |
| --- | --- | --- | --- | --- | --- | --- | --- | --- |
|  |  |  |  | Unadjusted for cage |  |  |  |  |
| Treatment | N | Median lifespan | % difference, to median lifespan Control | log-rank p-value | Cox PH log-rank p-value | 90<sup>th</sup> percentile lifespan | % difference, to 90<sup>th</sup> percentile Control | Wang-Allison p-value |
| *Control* | 311 | 792 | - | Ref. | Ref. | 1079 | - | Ref. |
| *Asta* | 156 | 908 | 14 | **0.003** | **0.003** | 1144 | 6 | **0.022** |
| *DMF_16* | 156 | 778 | -2 | 0.906 | 0.906 | 1078.5 | 0 | >0.999 |
| *DMF_early* | 156 | 813.5 | 3 | 0.265 | 0.265 | 1119 | 4 | 0.192 |
| *MEC* | 155 | 864 | 8 | **0.027** | **0.027** | 1138.8 | 5 | 0.102 |
| *MPA* | 156 | 808 | 2 | 0.081 | 0.081 | 1139.5 | 6 | **0.050** |
| *PBA* | 155 | 832 | 5 | 0.186 | 0.186 | 1115.6 | 3 | 0.191 |
| *SG1002_2* | 156 | 784 | -1 | 0.495 | 0.495 | 1103.5 | 2 | 0.192 |

**Table 1b.**
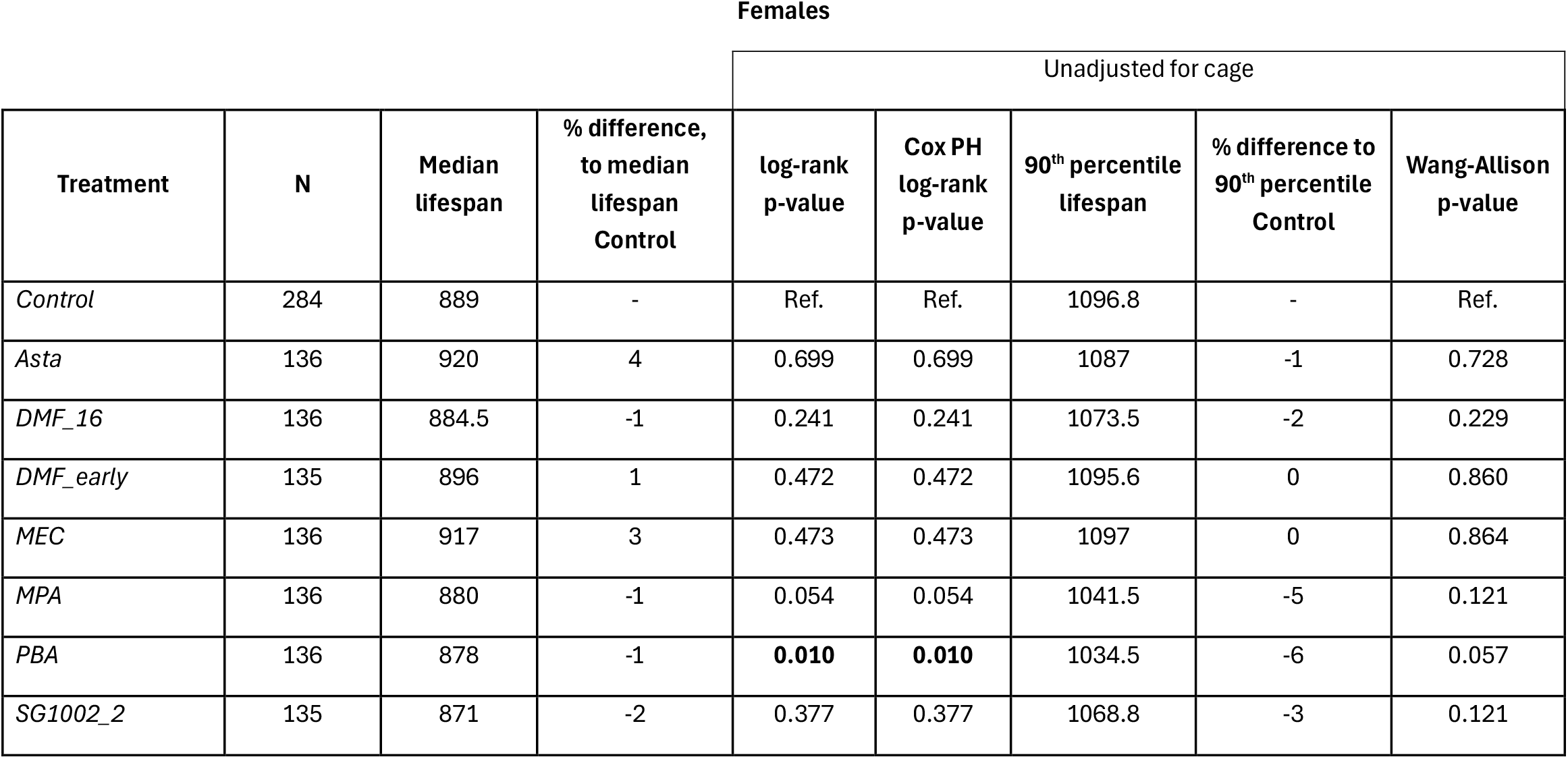
Standard lifespan data analyses, unadjusted for cage - for female animals only. Note that we have included p-values from both the log-rank test and unadjusted Cox models to illustrate that they are equivalent tests when unadjusted for random effects or additional covariates (*Stefanescu, 2003;* Harrell & Harrell, 2015).

For female mice, we saw that only animals treated with *PBA* showed significantly lower median survival time as compared with control (879 vs 893 days, *p* = 0.010), along with lower maximum lifespan which was not statistically significant at the 0.05 level (1034.5 vs 1096.8 days, *p* = 0.057; Table 1b).

### Linear Models Adjusted vs. Unadjusted for Cage and Nesting effects

Linear mixed models examining the effect of treatment and sex on the longevity of non-censored subjects displayed similar results to the unadjusted log-rank tests overall in terms of nominal significance (p<0.05), and particularly for male mice (Table 2a). In males, *Asta* and *MEC* treated animals survived significantly longer than control animals, as in the results seen in the simple log-rank tests (857/843 days vs 773 days, respectively; Table 2a). Female mice showed no significant differences in survival between treatments (Table 2b), and thus do not show the same significant reduction in survival from *PBA* treatment observed in unadjusted models (Table 1b).

**Table 2.** Linear model results on longevity of non-censored animals. Models accounting for correlation within cages (“adjusted”) use linear mixed models with random effects for cage and cage nested within site, and degrees of freedom calculated using the Kenward-Roger approximation method. Note that the p-values of the type 3 test were found to be identical between SAS and R. Models include all animals with factors for treatment, sex, and treatment x sex interaction.

| Type 3 tests | Unadjusted for Cage, Linear Models |  |  | Adjusted for Cage, Linear Mixed Models |  |  |
| --- | --- | --- | --- | --- | --- | --- |
|  | F | Num DF, Den DF | p-value | F | Num DF, Den DF | p-value |
| Treatment | 3.23 | 7, 2553 | **0.002** | 2.64 | 7, 659 | **0.011** |
| Sex | 54.80 | 1, 2553 | **<0.001** | 44.74 | 1, 659 | **<0.001** |
| Treatment * Sex | 1.35 | 7, 2553 | 0.221 | 1.13 | 7, 659 | 0.340 |

**Table 2a.** Estimated means and standard error for each treatment group in male animals. P-values for estimated means are from pairwise comparisons of each treatment mean vs. the control.

| Males |  |  |  |  |  |  |
| --- | --- | --- | --- | --- | --- | --- |
|  | <u>Unadjusted for Cage, Linear Models</u> |  |  | <u>Adjusted for Cage, Linear Mixed Models</u> |  |  |
| Treatment | Estimated Means | SE | P-value | Estimated Means | SE | P-value |
| *Control* | 773 | 12.7 | NA | 773 | 18.9 | NA |
| *Asta* | 857 | 17.5 | **< 0.001** | 857 | 23.0 | **< 0.001** |
| *DMF_16* | 776 | 17.9 | 0.895 | 775 | 23.4 | 0.923 |
| *DMF_early* | 814 | 17.9 | 0.062 | 814 | 23.3 | 0.081 |
| *MEC* | 843 | 17.9 | **0.001** | 844 | 23.4 | **0.003** |
| *MPA* | 814 | 17.9 | 0.061 | 814 | 23.3 | 0.085 |
| *PBA* | 812 | 17.8 | 0.075 | 812 | 23.2 | 0.100 |
| *SG1002_2* | 775 | 17.8 | 0.919 | 775 | 23.3 | 0.926 |

**Table 2b.** Estimated means and standard error for each treatment group in female animals. P-values for estimated means are from pairwise comparisons of each treatment mean vs. the control.

| Females |  |  |  |  |  |  |
| --- | --- | --- | --- | --- | --- | --- |
|  | <u>Unadjusted for Cage, Linear Models</u> |  |  | <u>Adjusted for Cage, Linear Mixed Models</u> |  |  |
| Treatment | Estimated Means | SE | P-value | Estimated Means | SE | P-value |
| *Control* | 881 | 13.0 | NA | 881 | 19.0 | NA |
| *Asta* | 904 | 19.0 | 0.315 | 904 | 25.0 | 0.390 |
| *DMF_16* | 859 | 18.7 | 0.326 | 859 | 24.7 | 0.374 |
| *DMF_early* | 859 | 18.8 | 0.338 | 859 | 24.8 | 0.382 |
| *MEC* | 897 | 18.7 | 0.483 | 897 | 24.7 | 0.538 |
| *MPA* | 860 | 18.9 | 0.368 | 860 | 24.9 | 0.416 |
| *PBA* | 857 | 18.8 | 0.297 | 857 | 24.7 | 0.348 |
| *SG1002_2* | 871 | 18.9 | 0.649 | 871 | 24.9 | 0.683 |

When results from these mixed models accounting for cage clusters and nesting were compared to those from standard linear models that did not account for clustering and nesting, we see that the standard errors (SE) of the estimates are larger (as expected after accounting for clustered data), and that the denominator degrees of freedom are on the order of the number of cages (n = 761), which resulted in small changes in p-values. However, no difference in statistical conclusions (*p*<.05) were detected in this case-study for the male or female mice (for males: *p* < 0.001 for *Asta* for both adjusted and unadjusted, and *p* = 0.001 *and p* = 0.003 for *MEC* for unadjusted and adjusted respectively; Table 2a).

### Full Cox Frailty Models Adjusted for Cage and Nesting effects

Finally, Cox PH and Cox frailty models fitted with both Gaussian and gamma distributions for the random cage and site effects and including all treatment groups, sex, and an interaction between treatment and sex showed similar overall results to those seen from the linear mixed models (Table 3). Specifically, we found that male mice in the *Asta* (HR = 0.69 / HR = 0.68, *p* = 0.001 for gamma/Gaussian models; Table 3a) and *MEC* (HR = 0.75, *p* = 0.016 / HR = 0.75, *p* = 0.019 for gamma/Gaussian models; Table 3a) groups showed significantly reduced death hazard as compared to control, indicating longer survival time. For females, no treatment groups showed significant changes in death hazard as compared to the control group in frailty models (Table 3b), like was seen in the linear mixed model results.

**Table 3.** Cox PH and frailty model results on longevity. These “full” models include variables for treatment, sex, and an interaction between treatment and sex, and (in the case of the frailty models) a frailty term for cage nested within site.

| Type 3 tests | Unadjusted, Cox PH |  | Gamma distribution of frailty term |  | Gaussian distribution of frailty term |  |
| --- | --- | --- | --- | --- | --- | --- |
|  | LR Chi-square | p-value | Wald Chi-square | p-value | Wald Chi-square | p-value |
| Treatment | 16.31 | **0.022** | 14.77 | **0.013** | 13.85 | **0.014** |
| Sex | 0.28 | 0.600 | 1.49 | 0.158 | 1.53 | 0.142 |
| Treatment * Sex | 12.63 | 0.082 | 8.34 | 0.149 | 8.10 | 0.137 |

**Table 3a.** Estimated hazard ratios between each treatment group and the control for male animals. P-values are for the significance of each pairwise treatment comparison vs. the control. Standard errors reported are on the log hazards scale.

|  | Males |  |  |  |  |  |  |  |  |
| --- | --- | --- | --- | --- | --- | --- | --- | --- | --- |
|  | Unadjusted, Cox PH |  |  | Gamma distribution of frailty term |  |  | Gaussian distribution of frailty term |  |  |
| Treatment | Estimated HR | SE | P-value | Estimated HR | SE | P-value | Estimated HR | SE | P-value |
| *Control* | 1.00 | NA | NA | 1.00 | NA | NA | 1.00 | NA | NA |
| *Asta* | 0.71 | 0.10 | **< 0.001** | 0.69 | 0.12 | **0.001** | 0.68 | 0.12 | **0.001** |
| *DMF_16* | 1.01 | 0.10 | 0.924 | 1.01 | 0.12 | 0.924 | 1.01 | 0.12 | 0.914 |
| *DMF_early* | 0.87 | 0.10 | 0.171 | 0.87 | 0.12 | 0.240 | 0.87 | 0.12 | 0.254 |
| *MEC* | 0.76 | 0.10 | **0.007** | 0.75 | 0.12 | **0.016** | 0.75 | 0.12 | **0.019** |
| *MPA* | 0.79 | 0.10 | **0.021** | 0.80 | 0.12 | 0.061 | 0.80 | 0.12 | 0.067 |
| *PBA* | 0.85 | 0.10 | 0.107 | 0.87 | 0.12 | 0.242 | 0.87 | 0.12 | 0.239 |
| *SG1002_2* | 0.88 | 0.10 | 0.194 | 0.91 | 0.12 | 0.451 | 0.91 | 0.12 | 0.450 |

**Table 3b.**
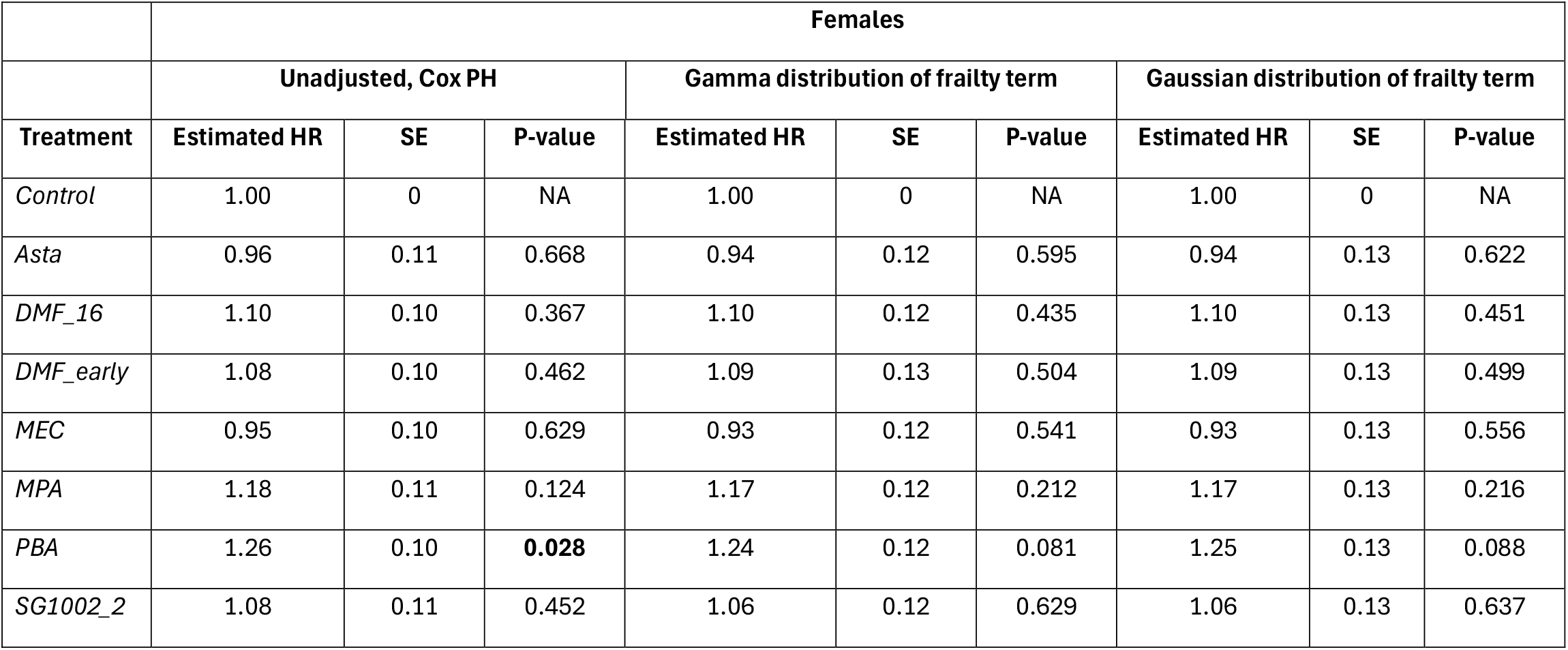
Estimated hazard ratios between each treatment group and the control for female animals. P-values are for the significance of each pairwise treatment comparison vs. the control. Standard errors reported are on the log hazards scale.

|  | **Females** |  |  |  |  |  |  |  |  |
| --- | --- | --- | --- | --- | --- | --- | --- | --- | --- |
|  | **Unadjusted, Cox PH** |  |  | **Gamma distribution of frailty term** |  |  | **Gaussian distribution of frailty term** |  |  |
| **Treatment** | **Estimated HR** | **SE** | **P-value** | **Estimated HR** | **SE** | **P-value** | **Estimated HR** | **SE** | **P-value** |
| *Control* | 1.00 | 0 | NA | 1.00 | 0 | NA | 1.00 | 0 | NA |
| *Asta* | 0.96 | 0.11 | 0.668 | 0.94 | 0.12 | 0.595 | 0.94 | 0.13 | 0.622 |
| *DMF_16* | 1.10 | 0.10 | 0.367 | 1.10 | 0.12 | 0.435 | 1.10 | 0.13 | 0.451 |
| *DMF_early* | 1.08 | 0.10 | 0.462 | 1.09 | 0.13 | 0.504 | 1.09 | 0.13 | 0.499 |
| *MEC* | 0.95 | 0.10 | 0.629 | 0.93 | 0.12 | 0.541 | 0.93 | 0.13 | 0.556 |
| *MPA* | 1.18 | 0.11 | 0.124 | 1.17 | 0.12 | 0.212 | 1.17 | 0.13 | 0.216 |
| *PBA* | 1.26 | 0.10 | **0.028** | 1.24 | 0.12 | 0.081 | 1.25 | 0.13 | 0.088 |
| *SG1002_2* | 1.08 | 0.11 | 0.452 | 1.06 | 0.12 | 0.629 | 1.06 | 0.13 | 0.637 |

When comparing the Cox PH models adjusted for clustering and nesting to the unadjusted Cox PH models, we see that the SEs and p-values are both a bit larger. For both the males and females, the unadjusted Cox PH model showed significant effects (Males: MPA, HR=0.79, *p*=0.021; Females: PBA, HR=1.26, *p*=0.028) that were not maintained in the adjusted models.

In general, the Cox PH models unadjusted for cage and nesting effects showed artificially deflated standard errors and p-values as compared to the models which properly accounted for the known cage and nesting effects.

## Discussion

The present study investigated the impact of clustering and nesting effects on lifespan analyses using data from the Interventions Testing Program (ITP). Our findings, particularly in the comparison between the Cox PH and Cox frailty models underscore the critical importance of accounting for clustering and nesting in preclinical aging research to ensure valid statistical inferences, even when the number of clusters is large.

### Key Findings and Implications

Our analysis revealed that, results from analyses using standard methods compared to results from analyses which properly accounted for cage effects showed some differences in statistical significance for both male and female mice. The use of linear mixed models and especially Cox frailty models, which account for cage-level dependencies, provided presumably more accurate (and larger) variance and p-value estimates when compared to unadjusted models, as expected from prior research on this topic (**Murray et al., 2004**; **Golzarri-Arroyo et al., 2022**). For these data we observe that if we had used only Cox PH models, we would have incorrectly concluded that *MPA* showed a significant treatment effect in males (Table 3a), and that *PBA* showed a significant treatment effect in females (Tables 1b and 3b) – effects which disappeared when cage effects were appropriately accounted for. These results highlight the critical need for researchers to incorporate clustering and nesting adjustments in their analyses, regardless of whether they expect such adjustments to change their conclusions. Ignoring clustering and nesting effects can lead to overestimated precision and increased risk of Type I errors, such as those seen here, potentially compromising the validity of study conclusions (**Pearl, 2014**). This is particularly relevant in aging research and animal studies, where animals are often housed in a relatively small number of shared cages, introducing dependencies that can distort variance estimates if not appropriately accounted for (**Leyrat et al., 2018; Li & Redden, 2015**).

### Methodological Considerations

The importance of accounting for clustering and nesting extends beyond the specific context of this study. In preclinical research, including aging studies, clustered designs are commonly used due to practical and ethical considerations in animal housing. However, the statistical implications of these designs are often overlooked, leading to potential biases in results and interpretations. Recognizing these challenges has driven the development of advanced techniques to account for complex designs (**Schnürch et al., 2021**).

Our study demonstrates the practical application of clustering and nesting adjustments using widely available statistical methods such as linear mixed models and Cox frailty models (**Austin, 2017; Balan & Putter, 2020**). These approaches allow researchers to model dependencies within clusters, ensuring that statistical conclusions accurately reflect the true structure of the data. By providing a framework applicable to other preclinical studies, our work contributes to enhancing the reproducibility and rigor of research findings in aging studies and beyond.

### Comparison with Previous Research

Our findings align with previous research emphasizing the importance of accounting for clustering and nesting in experimental designs. Studies by **Murray et al**. (2004) and **Golzarri-Arroyo et al**. (2022) have shown that clustering and nesting can significantly impact variance estimates and statistical power. Our study extends this body of work by providing a detailed case study using ITP data, illustrating the practical implications of clustering and nesting adjustments in lifespan analyses.

Furthermore, our results are consistent with broader trends in biomedical research calling for increased methodological rigor. For example, recent work by de **Jong et al**. (2020), **Chusyd et al**. (2022), and **Klatt et al**. (2023) has highlighted the persistent issue of overlooking clustering and nesting effects in preclinical studies, underscoring the need for greater awareness and implementation of appropriate statistical techniques.

Importantly, a recent study by **Luciano and Churchill** (2025) specifically examined the impact of co-housing on murine aging studies. Using a simulation-based approach and analysis methods similar to what we have done here, these authors showed that accounting for clustering and nesting effects is critical for valid inferences, and also highlight the importance of carefully considering housing conditions and their statistical implications in aging studies.

### Limitations and Future Directions

While our study provides valuable insights into the importance of accounting for clustering and nesting in animals studies generally, it is important to acknowledge its limitations. Future research should explore the consequences of not appropriately accounting for clustering and nesting in other datasets and experimental conditions in aging research. Additionally, further investigation is needed to refine clustering and nesting adjustment techniques and assess their impact on different types of preclinical outcomes. This could include the development of statistical methods for estimating effects on maximum lifespan while accounting for clustering and nesting and investigating the impact of these adjustments on other endpoints relevant to aging research, such as healthspan measures.

## Conclusion

Our study highlights the critical importance of accounting for clustering and nesting effects in preclinical aging research. Incorporating clustering and nesting adjustments enhances the validity of results, ultimately contributing to more rigorous and reproducible science. We strongly advocate for the adoption of these practices in future studies to strengthen the overall quality of preclinical research in aging and related fields.

## Code availability

All analytical code produced in the present study will be uploaded to a public repository on OSF, and for now are available upon reasonable request to the authors.

